# Altered postnatal chromatin development in the nucleus accumbens primes lifelong stress sensitivity

**DOI:** 10.1101/2024.04.12.589272

**Authors:** Rebekah L. Rashford, Lisa Z. Fang, Michael DeBerardine, Hye Ji J. Kim, Laura Hirschfield, Ella Cervi, Mason R. Barrett, Jeremy M. Thompson, Meaghan C. Creed, Catherine Jensen Peña

## Abstract

Early-life stress (ELS) sensitizes individuals to subsequent stressors to increase lifetime risk for psychiatric disorders. Within the nucleus accumbens (NAc) — a key limbic and reward-associated brain region — ELS sensitizes both cellular and transcriptional response to later stress, which are programmed by enduring epigenetic changes. Among the histone modifications persistently enriched by ELS in NAc is H3K4me1, which is associated with open chromatin and epigenetic priming of genomic enhancers. Here, we sought to determine whether H3K4me1 enrichment in NAc was sufficient to prime cellular and behavioral responses to adult stress. Viral-mediated overexpression of the histone H3 monomethyltransferase *Setd7* in juvenile NAc induced lifelong chromatin changes and predominately opened chromatin at long-range cis-regulatory elements predicted to enhance immediate early-genes and transcriptional regulators of mesolimbic development and synaptic activity. These epigenetic changes altered physiological properties of D2-type medium spiny neurons in NAc to resemble neurons of stressed mice, without significantly altering D1-type neurons. Finally, juvenile — but not adult — *Setd7* overexpression and H3K4me1 enrichment in NAc enhanced behavioral sensitivity to future stress. Together, these data indicate that altered postnatal chromatin development in NAc by H3K4me1 enrichment is sufficient to prime lifelong transcriptional, physiological, and behavioral stress sensitivity.

## INTRODUCTION

The neural epigenome continues to mature after birth and its maturation is fine-tuned by the environment (Peña, 2025; Stroud et al., 2020). Even after cell-fate specification, maturation of chromatin and DNA methylation patterns shape cell-type identity and function, with significant maturation occurring at enhancers — cis-regulatory regions of the genome that can act on genes many kilobases away (Long et al., 2016; Bonifer and Cockerill, 2017). Epigenetic maturation of enhancers occurs until approximately the third postnatal week in mice and adolescence in humans, after which little further age-related change occurs (Franklin et al., 2025; Heffel et al., 2024; Herring et al., 2022; Lister et al., 2013; Simmons et al., 2013; Stroud et al., 2017, 2020). Developmental perturbations to the epigenome may become crystalized at this point, with persistent effects on gene regulation across the lifespan (Weaver et al., 2004; Peña et al., 2013; Stroud et al., 2020; Peña, 2025; Rashford et al., 2025).

Experience of early life stress (ELS) can alter maturation of the epigenome (Burns et al., 2018; Peña, 2025; Rahman and McGowan, 2022; Geiger et al., 2024). Sequencing studies have found genome-wide alterations in DNA methylation and histone modifications in peripheral tissues and brain samples from individuals with a history of child maltreatment (Labonté et al., 2012; Lutz et al., 2021, 2017; Parade et al., 2021; Weder et al., 2014). These epigenetic alterations likely mediate persistent genome-wide changes in gene expression throughout the brain following ELS (Kos et al., 2023; Parel and Peña, 2020; Peña et al., 2019, 2017; Short et al., 2023). In turn, these molecular changes are thought to contribute to altered structural maturation of the brain, functional responses to cognitive tasks, stressors, and rewards, heightened stress response, and mediate increased risk for psychiatric disease observed following ELS (Hanson et al., 2021; Peña, 2025). However, it is not well understood whether altered maturation of the epigenome contributes to greater stress sensitivity and psychiatric disease risk compared to epigenetic changes that may accumulate in adulthood.

Dynamic changes to post-translational histone modifications have recently been identified in the nucleus accumbens (NAc) of mice exposed to ELS (Kronman et al., 2021). The NAc is a limbic brain region extensively implicated in the pathophysiology of psychiatric disease (Hanson et al., 2021). At a cellular level, ensembles of NAc neurons that are initially activated by experience of ELS are hyper-reactive to stress across the lifespan, contributing to stress sensitivity in mice (Balouek et al., 2023). At a transcriptional level, ELS induces long-lasting changes to baseline gene expression in NAc that are relevant for predicting successful antidepressant treatment response versus failure (Parel et al., 2023). Secondarily, ELS induces latent gene expression changes revealed by a second hit of stress in adulthood (Peña et al., 2019), pointing to epigenetic regulatory mechanisms at gene enhancers that may prime latent transcriptional responses (Geiger et al., 2024). Epigenetic priming is a form of molecular memory in which developmental or environmental stimuli open chromatin at enhancers to facilitate response to future stimuli, and which is marked by histone-3 lysine-4 monomethylation (H3K4me1) (Calo and Wysocka, 2013). We recently found that H3K4me1 was enriched in ventral tegmental area following ELS (Geiger et al., 2024), suggesting similar biological mechanisms may drive concerted epigenetic adaptations across limbic brain regions. However, previous research in NAc has only focused on chromatin modifications associated with active gene expression and adult manipulations (Covington et al., 2011; Heller et al., 2014; Kronman et al., 2021; Lepack et al., 2016; Sun et al., 2015; Torres-Berrío et al., 2024), leaving the role of developmental priming completely uncharacterized.

Here, we sought to determine whether developmental alterations in the NAc epigenome following ELS are sufficient to prime heightened stress response. To test this, we used a combination of NAc histone bottom-up mass spectrometry, overexpression of the H3K4me1-specific methyltransferase *Setd7*, ATAC-sequencing, patch-clamp physiology in D1- and D2-type medium spiny neurons, and characterization of behavior before and after sub-chronic adult social defeat stress. Finally, we tested whether NAc enrichment of H3K4me1 in postnatal development versus adulthood uniquely sensitized stress-related behaviors. Our results provide novel insights into how altered development of the epigenome may encode long-lasting stress sensitivity.

## RESULTS

### Enriched H3K4me1 following ELS

To determine whether ELS altered development of chromatin modifications associated with enhancer priming and activity in NAc, we took advantage of published bottom-up mass spectrometry data that profiled more than 200 individual and combinatorial post-translational histone modifications in the NAc at P21, P35, and P70-80 following ELS (Kronman et al., 2021). We found both a main effect of age (two-way ANOVA, F(2,12)=42.86, *p*<0.0001), and a main effect of ELS on H3K4me1 (F(2,12)=4.759, *p*=0.049), such that after weaning both aging and ELS increased H3K4me1 (**Figure 1A**). This early enrichment suggests that ELS augments age-related enrichment of H3K4me1.

**Figure 1.**
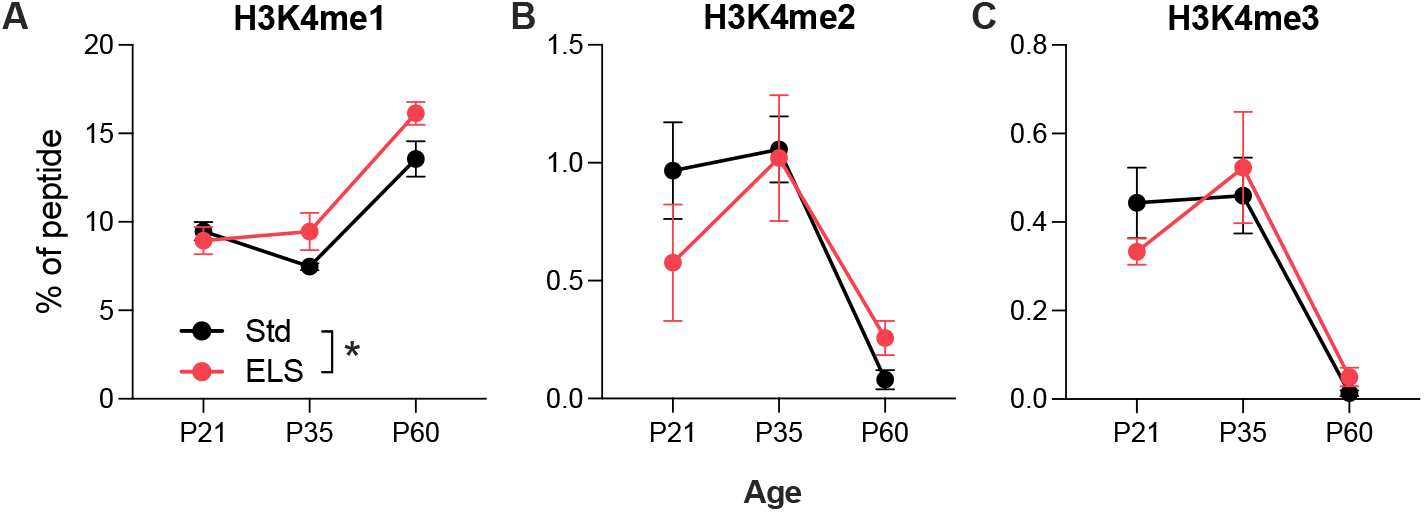
ELS selectively enriches for H3K4me1. Proportion of (**A**) H3K4me1, (**B**) H3K4me2, and (**C**) H3K4me3 peptide fragments in male NAc across postnatal development (reanalyzed mass spectrometry data published in (Kronman et al., 2021). Mean±SEM; \**p*<0.05 for main effect of ELS.

In contrast, di- and tri-methylation at H3K4 — histone modifications associated with active gene expression — decreased with age [main effect of age for H3K4me2: F(2,12)=11.87, *p*=0.0014; main effect of age for H3K4me3 F(2,12)=22.69, *p*<0.0001], but were unaffected by ELS (**Figures 1B-C**). Active enhancers are further distinguished from primed enhancers by the presence of H3K27Ac. Although this modification was below detection levels at P21 and P35, at P60, H3K27Ac was enriched in standard-reared NAc but not ELS (Welch’s two-tailed t=14.50, *p*=0.0002; **Supplemental figure S1A**). We also found an interaction between ELS and age for H3K27me2, a histone modification associated with protecting enhancers from spurious activation (Ferrari et al., 2014), such that this modification was decreased following ELS but elevated in adulthood (**Supplemental Figure S1B**; F(2, 12)=5.874, *p*=0.017). There were no main effects or interactions with ELS for H3K27me3 or H3K36me3, both associated with inhibition of gene expression (**Supplemental Figures S1C-D**). Taken together, our findings that ELS augments and accelerates H3K4me1 accumulation in the NAc without simultaneous changes in markers of active gene expression are consistent with both premature development and priming of chromatin architecture.

### Juvenile Setd7 overexpression alters lifelong chromatin accessibility in NAc

Given the specific enrichment of H3K4me1 following ELS and the role of H3K4me1 in priming enhancers (Griffith et al., 2024; Heintzman et al., 2007; Mercer et al., 2011), we sought to test the impact of early accumulation of H3K4me1 in NAc on chromatin accessibility state. To do this, we took advantage of an epigenome editing tool recently validated by our group to over-express the histone mono-methyltransferase *Setd7* to enrich for H3K4me1 (Geiger et al., 2024). AAV-*Setd7* broadly enriches for H3K4me1 without inducing further H3K4 methylation or other histone modifications associated with active gene expression, such as H3K27Ac. We hypothesized that juvenile *Setd7* overexpression and consequent H3K4me1 enrichment would induce a long-lasting open and accessible chromatin state associated with epigenetic priming. To test whether *Setd7* overexpression opened chromatin, we bilaterally overexpressed AAV-*Setd7* or AAV-*Gfp* (**Figure 2A**) in juvenile NAc at postnatal day P9-12 (*n*=5-6/group) and collected tissue punches for ATAC-sequencing in adulthood (P70-80).

**Figure 2.**
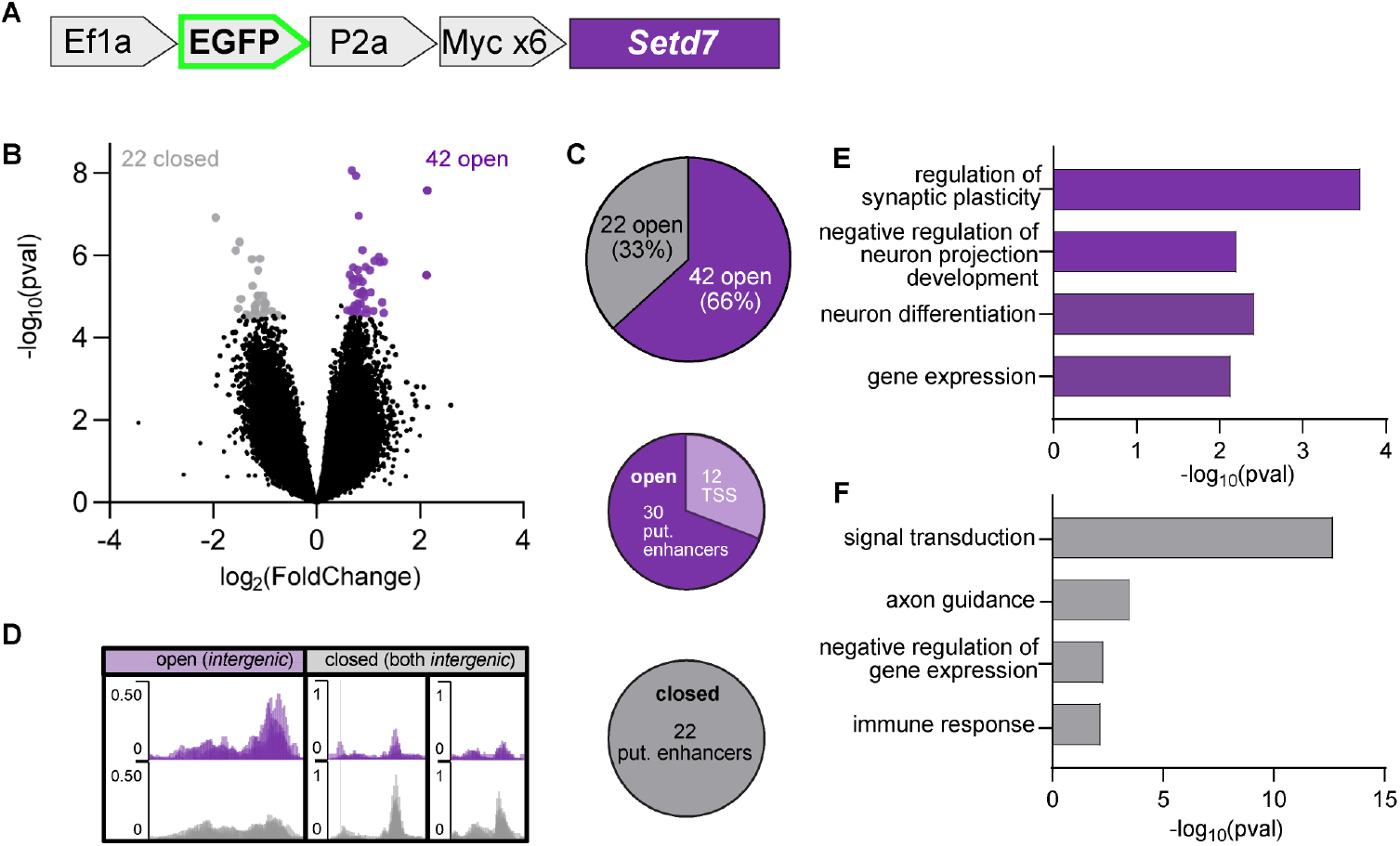
Juvenile *Setd7* overexpression alters lifelong chromatin accessibility in NAc. (**A**) Viral schematic of AAV9-EF1a-EGFP-P2A-MYC-Setd7; control vector was AAV9-EF1a-EGFP. (**B**) Volcano plot of bulk ATAC-seq showing chromatin regions with gained or lost accessibility with juvenile *Setd7* (*n*=5) vs. *Gfp* (*n*=6) overexpression. Significant genomic regions (p.adj.<0.1 and |log_2_FC| >0.5) are shown in purple (more open with *Setd7*) and gray (more closed). (**C**) Percentage of DARs opened vs closed (top). Among open (middle) and closed (bottom) DARs, proportion mapping to long-range cis-regulatory elements (putative enhancers) or to promoter regions. (**D**) Example RPM-normalized track overlays of an open region (left [chr10:59072341-59072625]) and closed regions (center [chr18-86218908-86219130], and right [chr4:27247388-27247650]). (**E**) Gene ontology enrichment for genes within or nearest to opened DARs. (**F**) Gene ontology enrichment for genes within or nearest to closed DARs.

Analysis of differentially accessible regions (DARs) showed that juvenile *Setd7* overexpression and H3K4me1 enrichment induced long-lasting changes in chromatin accessibility relative to *Gfp* controls, with nearly twice as many regions of opened vs closed chromatin (DESeq2 p.adj <0.1, |log_2_FC| >0.5; **Figure 2B-C; Supplemental Table 1**). Most differentially open chromatin regions and all closed chromatin regions were long-range cis-regulatory regions and putative enhancers (>1kb away from a transcription start-site and not within an exon; **Figure 2C**). Examples of both opened and closed intergenic regions are shown in **Figure 2D**. To determine the genes these DARs may be regulating (as enhancers do not always regulate their nearest gene neighbors; (Schoenfelder and Fraser, 2019)), we used a long-range target gene predictor tool based on publicly available histone modification and gene expression data (Grant and Bailey, 2021). Putatively primed genes (genes predicted to be regulated by open DARs) include immediate early genes and transcriptional regulators important for mesolimbic development and synaptic activity (*Fos, Homer1, Gadd45g, Cebpb, Cdkn1c, Prrx1*, and *Eif4ebp2*), genes encoding transmembrane receptors and channels (*Kcnq3, Grin1, Gab2*), and epigenetic regulators (*Tet1*) (**Supplemental Table 2**). Gene ontology (GO) analysis identified an enrichment of genes related to synaptic plasticity, neuron development and differentiation, and gene expression (**Figure 2E**). Putatively repressed genes (regulated by closed DARs) are enriched for those regulating various homeostatic processes, signal transduction, and axon guidance, among other processes (**Figure 2F, Supplemental Table 2**). Together, these results suggest developmental enrichment of H3K4me1 through *Setd7* overexpression in the NAc may facilitate response to future stimuli by driving a more open chromatin landscape and priming genes that promote transcriptional and synaptic activity.

### Juvenile Setd7 overexpression in NAc mimics stress phenotype in D2-MSNs

A majority of neurons of the NAc are medium spiny neurons (MSNs) that can be distinguished by their expression of either type 1 (*Drd1*; D1) or type 2 (*Drd2*; D2) dopamine receptors. D1 and D2-MSNs play opposing roles in mediating stress-relevant avoidance and appetitive behavior, with stimulation of D2-MSNs exerting lateral inhibition over D1-MSNs and generally promoting avoidance (Dobbs et al., 2016; Fang and Creed, 2024; Francis et al., 2014; Kravitz et al., 2012; Lemos et al., 2016; Lobo et al., 2010; Peña, 2017). To test the hypothesis that juvenile H3K4me1 enrichment via *Setd7* overexpression may alter neuronal physiology, we overexpressed *Setd7* in juvenile (P9-12) Drd1a-tdTomato mice and performed patch-clamp electrophysiology in D1- and D2-MSNs, distinguished based on presence (D1-MSNs) or absence of red fluorescence (D2-MSNs) (Creed et al., 2015; Pascoli et al., 2014). Putative D2-MSNs were also confirmed by absence of spontaneous or burst firing which may indicate interneurons ((Johansson and Silberberg, 2020; Kawaguchi, 1993), see methods).

We found no change in the excitability metrics of D1-MSNs (**Supplemental Figure S2A-G**) or their passive membrane properties (**Supplemental Figure S2A-G**) induced by *Setd7* virus expression or subchronic adult stress. In contrast, *Setd7* overexpression in D2-MSNs resulted in a leftward shift in the input-output curve of spikes per current injection [**Figure 3A;** interaction between current injection and *Setd7*: (F(30,600)=3.170, *p*<0.0001, two-way RM ANOVA], indicating subtle hyperexcitability of D2-MSNs. While D2-MSNs from GFP-injected mice that had undergone the subthreshold chronic defeat paradigm exhibited a similar leftward shift in the IO curve relative to unstressed GFP-controls, there was no additional effect of *Setd7* expression [**Figure 3B**; F(1,24)=3.806, *p*=0.0628, two-way RM ANOVA]. Together, this suggests that *Setd7* mimics the effect of subthreshold stress on current firing of D2-MSNs, with further occlusion of the *Setd7* effect in stressed mice. Bolstering this interpretation, the input resistance of D2-MSNs was significantly increased by *Setd7* overexpression (*p*=0.006, U=53.00; Mann-Whitney test), an effect which was again occluded following subchronic stress (**Figure 3D**; *p=*0.4184, U=68.00). There was no change in rheobase (Control: *p=*0.2219, *U*=69.00; Stress: *p*=0.4023, *U*=91.50, Mann-Whitney test) or resting membrane potential (**Figure 3C, E;** Control: *p*=0.2577, *U*=93.50; Stress: *p*=0.6498, *U*=75.00, Mann-Whitney test). Finally, *Setd7* overexpression increased the voltage sag elicited by hyperpolarizing current injections [**Figure 3F;** *F*(1,30)=5.039, *p*=0.0323, two-way RM ANOVA], an effect which was again mimicked by subthreshold defeat, with no additive effect of *Setd7* overexpression [**Figure 3G**; *F*(1,24)=0.6277, *p*=0.4359, two-way RM ANOVA. Given the opposing roles of D1- and D2-MSNs in mediating avoidance behavior, these findings lead to a prediction that *Setd7* overexpression in the NAc may subtly shift the balance between D1- and D2-MSNs towards promoting avoidance behavior.

**Figure 3.**
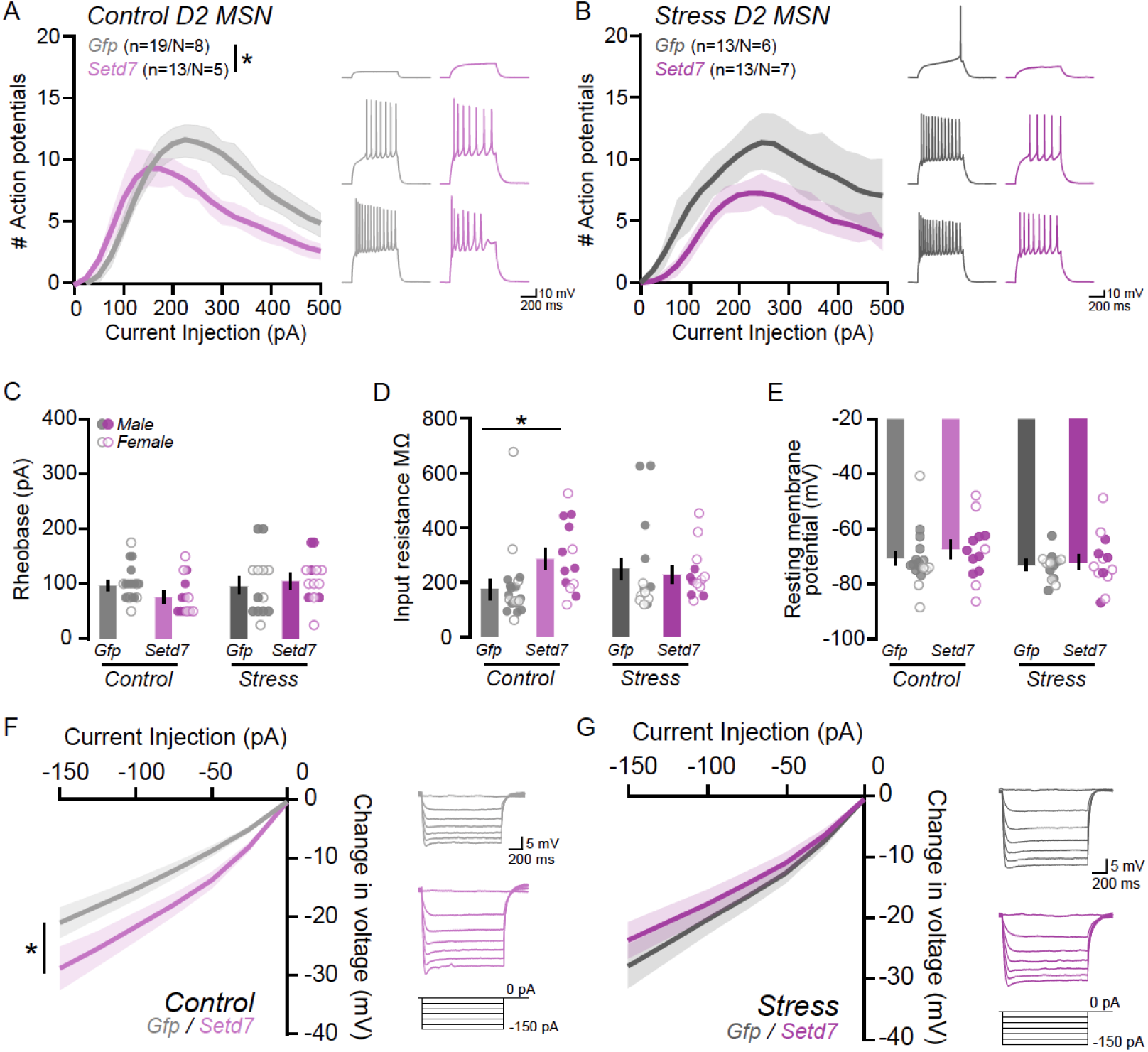
Juvenile *Setd7* overexpression increases the excitability of D2-MSNs, mimicking adulthood stress. **A-B**. (Left) Input-output plot of action potentials elicited in response to successive 600ms current injections from (**A**) control and (**B**) stressed mice. (Right) Representative traces depict action potential firing in response to a 50 pA (top), 150 pA (middle), and 250 pA (bottom) current injection. While there is no effect of Setd7 overexpression on action potential firing in D2 MSNs under, there is a significant interaction in control conditions, with a leftward shift in the IO curve. **C-E**. The effect of *Setd7* overexpression on intrinsic membrane properties, (**C**) rheobase, (**D**) input resistance, and (**E**) resting membrane potential, of D2-MSNs. *Setd7* overexpression does not alter the rheobase or resting membrane in either control or stressed mice but does increase the input resistance in control mice **F-G**. The peak change in voltage in response to successive 600ms hyperpolarizing currents from (**F**) control and (**G**) stressed mice. *Setd7* overexpression increases the peak voltage sag in control, but not stressed mice. \**p<*0.05; all data are depicted as mean ± SEM.

### Epigenetic priming via postnatal H3K4me1 enrichment in NAc increases sensitivity to future stress

We next tested whether developmental enrichment of H3K4me1 primes behavioral response to future stress. We again overexpressed AAV-*Setd7* or AAV-*Gfp* in NAc at P10 to match the start of ELS (**Figure 4A-B**; (Peña et al., 2017)). We used a within-subject design and mixed-effect modeling to test main effects and interactions of sex, virus, and stress (**Figure 4C**) on behavioral tests before and after five days of sub-chronic adult social defeat stress (Balouek et al., 2023; Francis et al., 2014; Yohn et al., 2019). Although we powered this study to detect sex differences (*n*=10-19/sex/group), there were no significant main effects of sex and results are presented together for simplicity. All statistical results are in **Supplemental Table 3**.

**Figure 4.**
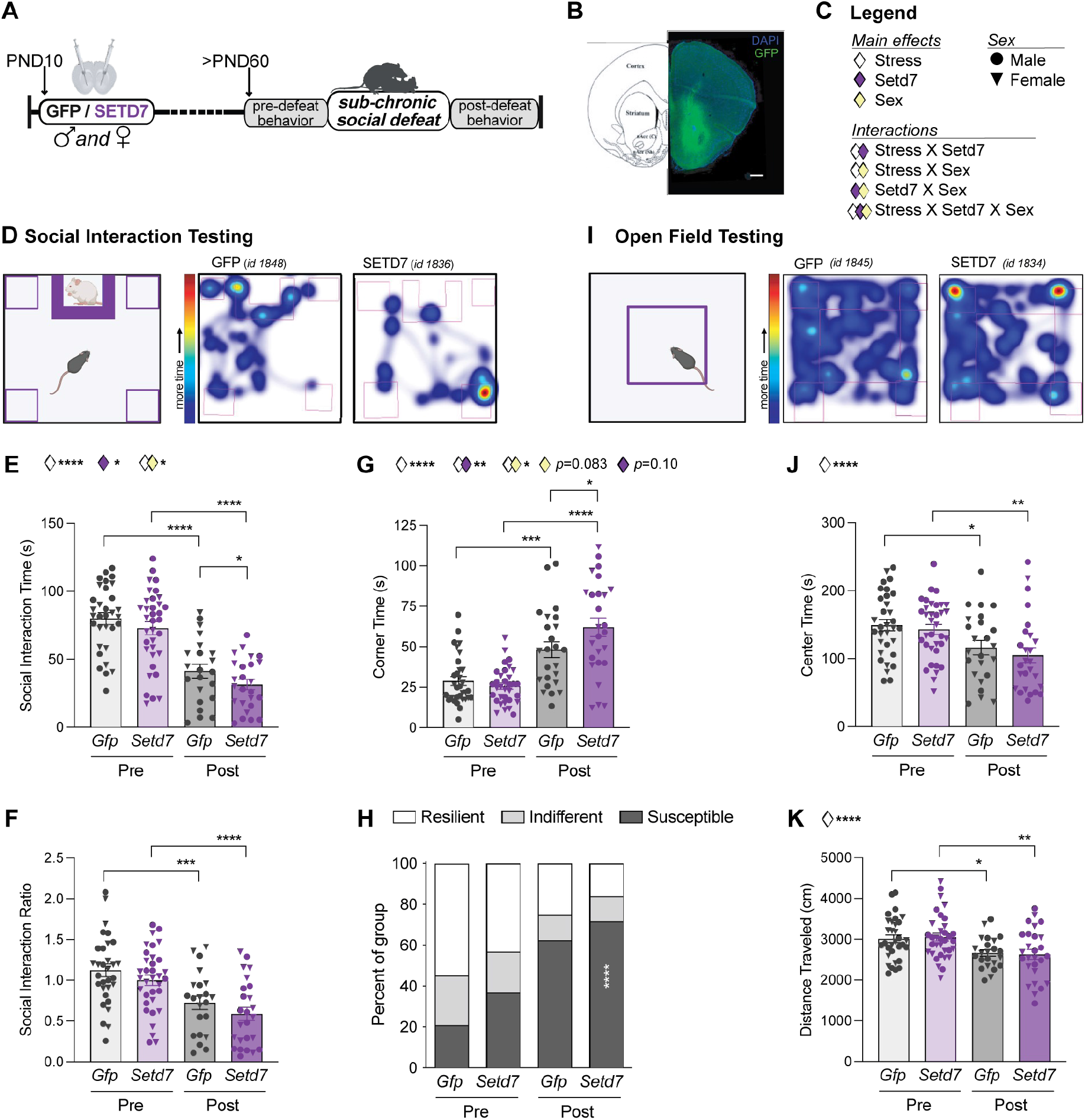
Juvenile *Setd7* overexpression in NAc increases behavioral sensitivity to future stress. (**A**) Experimental Timeline. (**B**) Representative image and brain atlas showing viral targeting of NAc. Scale bar, 1mm. (**C**) Legend for statistical main effects and interactions and individual male (circle) and female (triangle) data points. (**D**) Social interaction testing schematic: time spent in the interaction zone or corners was measured (left). Heatmaps showing representative trials from GFP and SETD7 groups, respectively (right). Within the social interaction test, (**E**) social interaction time, (**F**) social interaction ratio (lower ratio is associated with greater social avoidance), and (**G**) time spent in corners, among male and female mice before (pre) and after (post) social defeat stress. (**H**) Percent of each group considered resilient, indifferent, or susceptible based on SI ratio. (**I**) Open field testing schematic (left). Heatmaps showing representative trials from GFP and SETD7 groups, respectively (right). Within the open field test, (**J**) time spent in the center of the arena, and (**K**) total distance traveled. Main effects and interactions from mixed-effect modeling are indicated above each plot. Post hoc pairwise comparisons using Fisher’s LSD are depicted within each plot. \*\*\*\**p*<0.0001; \*\*\**p*<0.001; \*\**p*<0.01; \**p<*0.05; exact value displayed for trends *p*<0.1. Data are represented as mean ± SEM.

Across males and females, both stress and postnatal epigenetic priming by *Setd7* decreased social interaction time, a measure of depression-like behavior and stress sensitivity [**Figure 4D-E;** main effect of stress: F(1,44)=72.15, *p*<0.0001; main effect of *Setd7*: F(1,64) = 4.88, *p*=0.0306; interaction between stress and sex: F(1,44) = 6.54, *p*=0.014); no main effect of sex]. Similarly, both stress and *Setd7* decreased social interaction ratio [**Figure 4F;** main effect of stress: F(1,44)=20.61, *p*<0.0001; main effect of *Setd7*: F(1,64) = 5.148, *p*=0.027; trending interaction between stress and sex: F(1,44) = 2.959, *p*=0.0925)]. Time spent huddling in corners, an additional measure of heightened vigilance and stress sensitivity (Duque-Wilckens et al., 2020), was increased by stress in combination with *Setd7* and sex [**Figure 4G**; main effect of stress: F(1,44)=56.34, *p*<0.0001; interaction between stress and *Setd7*: F(1,44) = 7.705, *p*=0.008); interaction between stress and sex: F(1,44) = 4.229, *p*=0.046); trending main effect of sex: F(1,63) = 3.111, *p*=0.0826); trending main effect of *Setd7*: F(1,63) = 2.669, *p*=0.10)]. Collectively, *Setd7* and stress impacted the proportion mice categorized as susceptible, indifferent, and resilient based on social avoidance behavior (**Figure 4H**; *X*_2_=66.13, df=6, *p*<0.0001).

Time spent exploring the center of an open field (**Figure 4I**), a more generalized measure of threat avoidance, was sensitive to stress but not primed by juvenile *Setd7* overexpression [**Figure 4J**; main effect of stress: F(1,46)=22.93, *p*<0.0001; no main effects of *Setd7* or of sex]. Ambulatory behavior was not impaired by *Setd7* [**Figure 4K**; main effect of stress: F(1,42)=21.33, *p*<0.0001; no main effects of *Setd7* or of sex].

### Setd7 overexpression in adult NAc minimally influences stress response

We next sought to determine whether the impact of epigenetic priming was specific to developmental enrichment of H3K4me1, or whether *Setd7* overexpression in adulthood might similarly heighten sensitivity to later stress. We therefore expressed AAV-*Setd7* or AAV*-Gfp* in adult NAc (P60-70), waited three weeks, and tested behavior before and after sub-chronic social defeat stress as before (**Supplemental Figure S3A-C**). All statistical results are in **Supplemental Table 3**.

In contrast to juvenile *Setd7* overexpression, there was no main effect of stress but there was a main effect of adult *Setd7* expression to decrease social interaction time, but this effect did not further interact with stress [**Supplemental Figure S3D;** main effect of *Setd7*: F(1,30)=5.709, *p*=0.023; interaction between stress and sex: F(1,28) = 11.17, *p*=0.002; interaction between *Setd7* and sex: F(1,30) = 9.941, *p*=0.004)]. Adult *Setd7* overexpression also increased time spent in arena corners, a classic metric of avoidance behavior, but did not further interact with stress [**Supplemental Figure S3F;** main effect of *Setd7*: F(1,30)=4.022, *p*=0.050].There were no main effects of stress, *Setd7*, or sex on social interaction ratio (**Supplemental Figure S3E**), and the percent of mice categorized as susceptible was unchanged (**Supplemental Figure S3G**). Interestingly, adult *Setd7* modestly increased rather than decreased time spent exploring the center of an open field [**Supplemental Figure S3H;** main effect of *Setd7*: F(1,30)=4.178, *p*=0.050] but did not alter distance traveled (**Supplemental Figure S3I**). These results indicate that *Setd7* overexpression in adulthood is insufficient to increase sensitivity to subchronic social defeat stress. Taken together, juvenile — but not adult — *Setd7* overexpression and H3K4me1 enrichment in NAc heightens sensitivity to stress later in adulthood, suggesting postnatal development is particularly sensitive to epigenomic perturbations.

## DISCUSSION

Our findings presented here provide strong evidence that changes in the postnatal epigenome are carried forward across the lifespan and can impact neurophysiological and behavioral response to experiences such as stress. We show that ELS accelerates and augments postnatal accumulation of H3K4me1, and that postnatal enrichment of H3K4me1 through overexpression of the histone mono-methyltransferase *Setd7* alters physiological properties of D2-MSNs in the NAc and sensitizes behavioral response to adult social stress. These results help elucidate the mechanisms through which ELS programs long-lasting stress sensitivity and stress-induced risk for psychiatric disease through changes in molecular brain development.

Both acute and chronic stress dynamically alter post-translational histone modifications (Bastle and Maze, 2019; Covington et al., 2011; Farrelly et al., 2019; Geiger et al., 2024; Heller et al., 2014; Howerton et al., 2013; Kronman et al., 2021; Lepack et al., 2020, 2016; Levine et al., 2012; Morrison et al., 2020; Pusalkar et al., 2015; Rashford et al., 2025; Sun et al., 2015; Torres-Berrío et al., 2024). Through analysis of mass spectrometry data of histone modifications in NAc, we find that ELS augments levels of H3K4me1, a chromatin modification associated with epigenetic priming, without concomitant increases in histone modifications associated with active gene expression. ELS also increases H3K4me1 levels in VTA (Geiger et al., 2024) and drives open chromatin in ELS-activated cells of VTA (Rashford et al., 2025) suggesting epigenetic priming as a concerted response to ELS across limbic brain regions.

Examination of histone modifications across multiple ages also indicates that H3K4me1 accumulation may represent accelerated epigenetic development, given that levels of this histone modification increase across age. Similarly, ELS accelerated NAc accumulation of histone variant H3.3, associated with activity-dependent neuronal transcription, after a period of initial suppression (Lepack et al., 2016). ELS has also been found to accelerate other aspects of brain development, such as connectivity between amygdala and prefrontal cortical regions (Gee et al., 2013) and markers of hippocampal development including parvalbumin interneuron maturation, myelination, and synaptic plasticity (Bath et al., 2016). Accelerated development within some cell types or brain regions decouples otherwise tightly-controlled neurodevelopmental processes, which has been hypothesized to prematurely reduce plasticity and the ability to appropriately learn from expected environmental experiences (Rakesh et al., 2023; Reh et al., 2020). Our results support an interpretation that these effects may arise from accelerated epigenetic development.

Our findings intriguingly show that developmental accumulation of H3K4me1 in NAc, but not adult-specific accumulation of H3K4me1, is sufficient to prime heightened behavioral sensitivity to stress. This may be due to prolonged developmental maturation and plasticity of the epigenome through the postnatal juvenile period that finally crystalizes in adulthood (Franklin et al., 2025; Heffel et al., 2024; Herring et al., 2022; Lister et al., 2013; Simmons et al., 2013; Stroud et al., 2017, 2020). However, it has yet to be determined whether adult-specific *Setd7* overexpression also fails to broadly alter chromatin opening or whether perhaps a different set of genomic regions may be altered. Future studies may also test whether transient *Setd7* enzymatic activity in the juvenile period (as opposed to the long-term expression of AAVs) would become crystalized in the epigenome and carried forward to have similar persistent effects on physiology and behavior.

While *Setd7* overexpression here was not cell-type specific, prior single-nucleus RNA-seq within NAc shows that *Setd7* is expressed in nearly every cell type, with highest levels in D1 and D2 neuronal clusters and lowest levels in microglia and astrocytes (Ianov and Day, 2024; Phillips et al., 2023). Nevertheless, our results indicate a D2-specific neurophysiological mechanism. This is in line with prior work showing that repeated activation of D2-MSNs in stress-naïve mice induces sensitivity to later subthreshold social defeat stress (Francis et al., 2014). However, other studies have reported greater changes on D1-MSNs relative to D2-MSNs following chronic social defeat stress (Francis et al., 2014; Pignatelli et al., 2021), which we did not detect here. This may be due to the use of exclusively male mice in prior studies, owing to challenges in applying the chronic social defeat stress models in females. Moreover, prior changes in excitability were observed only in response to suprathreshold stress, with resilient individuals showing much more subtle physiological changes, consistent with the subtle adaptations we report following subchronic social defeat stress here. Finally, it is still unclear whether physiological adaptations on MSNs following chronic stress manipulations may reflect compensatory or homeostatic adaptations, since stimulating D1-MSNs promotes resilience, despite the finding of increased excitability associated with stress susceptibility (Francis et al., 2014; Francis and Lobo, 2016; Lobo et al., 2010). In this context, our results are consistent with a large body of prior work suggesting that the balance between D1- and D2-MSNs mediates approach and avoidance behavior, with increased D2-MSN activity relative to D1-MSN activity being predicted to promote avoidance behavior as we see here (Bariselli et al., 2019; Creed, 2018; Creed et al., 2016; Francis and Lobo, 2016; Kravitz et al., 2012; Matikainen-Ankney et al., 2023; Peña, 2017).

In sum, these data provide novel evidence that altered postnatal chromatin maturation is sufficient to prime sensitivity to future stress. Our research provides an important foundation for additional research to develop therapeutic strategies targeting chromatin to ameliorate the impact of ELS on brain development and psychiatric disease risk.

## Supporting information

Supplemental Table 1 Setd7 NAcATAC DARs

Supplemental Table 2 DAR-Primed genes

Supplemental Table 3 BehaviorStats

## ACKNOWLEDGEMENTS

Thank you to Hope Kronman for sharing raw mass-spec data published in (10.1038/s41593-021-00814-8). Thank you to Julie-Anne Balouek for generating and validating the Setd7 overexpression construct and for training and consultation on methods. Thank you to Shannon Bennett, Mayowa Oke, and Nibal Arzouni who contributed to an earlier version of this manuscript. We also thank Wei Wang and Jennifer Miller of the Princeton Genomics Core for conducting library preparation and sequencing. Parts of some figures were generated with BioRender.

## Funding

This research was funded by NIH K99MH115096, R00MH115096, NIH R01MH129643, New York Stem Cell Foundation(CJP);NIH5T32MH065214,NIHF31MH131351 (RR); R01DA049924, R01DA058755, R01DA056829 (MCC); Princeton Neuroscience Institute Newman Biousse Award (EC); Foundation for Anesthesia Education and Research Alison Cole endowed Mentored Research Training Grant (JMT); CIHR Postdoctoral Fellowship (LZF). CJP is a New York Stem Cell Foundation Robertson Investigator.

## AUTHOR CONTRIBUTIONS

The chromatin and behavioral studies were designed by CJP and RLR. ATAC-seq data were collected by RLR and analyzed by RLR and MD. Viral manipulations, behavioral work, and immunohistochemistry were performed by RLR, HJJK, LH, EC and MRB. The physiology experiments were designed by MCC with input from CJP, performed by LZF, JMT, and analyzed by LZF, JMT, and MCC. The manuscript was written by RLR, MCC, and CJP with input from all authors.

## COMPETING INTERESTS

There are no conflicts of interest to report.

## METHODS

### Mice

All experiments were conducted in accordance with the guidelines of the Institutional Animal Care and Use Committee at Princeton University (chromatin and behavioral experiments) and Washington University in St. Louis (electrophysiology experiments). Wild-type C57BL/6J mice were purchased (Jackson Laboratory) and housed in temperature- and humidity-controlled vivarium facilities and maintained on a 12-h light/dark cycle with *ad libitum* access to food and water. Breeding was performed in-house in trios; males were removed after 5-7 days, and females were separated into individual cages 1-3 days prior to parturition. Mice were considered male or female based on external genitalia (adult) and/or anogenital distance (pups). All offspring were weaned at postnatal day P21, with males and females weaned into separate cages, keeping littermates together and only combining pups from different litters of the same experimental condition in order to maintain 3-5 mice/cage. Males and females were used in all experiments. Retired Swiss-Webster male breeder mice (Taconic) were housed individually and used as aggressor mice in social defeat stress.

### Stereotaxic surgery

Standard-reared male and female C57BL/6J pups were randomly assigned to either experimental *Setd7* over-expression group or a control *Gfp* group. Surgeries were performed as previously described (Geiger et al., 2024) with the exception of NAc-targeting surgical coordinates. Adeno-associated viral vectors (AAV9) carrying a *Gfp*- and *Myc*-tagged *Setd7* transgene (pAAV-EF1a-EGFP-P2A-Myc-Setd7-WPRE-hGHpA) were previously developed (Geiger et al., 2024) from cDNA for *Setd7* (#87131; originally gifted from Francesca Spagnoli; (Kofent et al., 2016)) and *Gfp* (pAAV-EF1a-EGFP-WPRE-hGHpA) purchased from Addgene (#60058; originally gifted from Brandon Harvey; (Savolainen et al., 2014)). AAV-*Setd7* and AAV-*Gfp* control (identical, without *Myc-Setd7*) vectors were cloned and packaged by the Princeton Neuroscience Viral Core.

For juvenile overexpression, P9-12 pups were anesthetized with isoflurane (2-3% induction; 1-3% maintenance) and fitted into dual-arm stereotax with the nose fitted in the nose cone above the bite bar and the head stabilized using the blunt ends of ear bars between the jaw and ear. The bilateral surgical coordinates for pups were +/-1.4 (x = medial/lateral plane), +1.0(y = anterior/posterior plane), −3.15(z = dorsal/ventral plane), with both Hamilton syringes angled at 7°. The bilateral surgical coordinates for adults were +/-1.7 (x = medial/lateral plane), +1.6(y = anterior/posterior plane), −4.40(z = dorsal/ventral plane), with both Hamilton syringe angled at 10°. 250-300nL of virus was administered into each hemisphere at a rate of 0.1µL/minute. Both vet glue (Covetrus, cat#: 001505) and two-four sterile sutures (Fisher, cat#: 501180710) were used to ensure complete healing of incisions. Mice were given a perioperative topical dose of bupivacaine (0.25%, 2 mg/kg), and an additional dose 24h following surgery. Pups recovered in a clean cage on a warming pad before being returned to their home cage. To prevent cannibalization of pups by dams following surgery, pups were only returned to the home cage with the dam once completely ambulatory. All mice were monitored for five days post-surgery. Pups were weaned at P21 and aged to adulthood in standard group housing conditions. Adult mice that underwent surgery recovered for 3 weeks prior to stress, behavioral testing, and/or tissue collection. For molecular analysis and adult over-expression behavior experiments, viral targeting was assessed at the time of sacrifice by GFP flashlight (Electron Microscopy Sciences Nightsea flashlight, NC0792511). For juvenile over-expression behavior experiments, viral targeting was assessed by IHC (see below). In all experiments, mice with poor viral targeting were excluded from analysis.

### Sub-chronic social defeat stress

In order to determine whether chromatin manipulation increased sensitivity to later stress, male and female mice were subject to sub-chronic adult social defeat stress for 5 consecutive days. Previous studies have found that among males, susceptibility to social defeat stress develops across 10 days (chronic), but is not apparent after only 5 (Dias et al., 2014; Hodes et al., 2015; Muir et al., 2020). Here, both male and female mice were exposed to social defeat but were run in separate cohorts. Males were defeated as described in standard protocols (Golden et al., 2011). Females were defeated in a “non-discriminatory” version of social defeat as previously described (Bennett et al., 2024; Geiger et al., 2024; Yohn et al., 2019). Briefly, on each day of social defeat, experimental mice were placed into the cage of a novel Swiss-Webster retired breeder (“aggressor”; Taconic) mouse. Male lures were introduced first for ~1-2 minutes, then an experimental female mouse was added into the cage for an additional 5 minutes. After defeat, mice were moved across a perforated plexiglass barrier for the remainder of the day for continued sensory contact. Experimental mice were introduced to a new aggressor each day. Control mice were housed in a standard mouse cage in pairs of the same sex, separated by a perforated plexiglass barrier. Behavioral testing began one day after the last bout of social defeat.

### Behavioral paradigms and analysis

Mice were tested on a set of two behavioral tests twice for within-subject controls: once prior to (“pre”) and once after social defeat stress (“post”). Groups are presented as GFP-Pre (AAV-*Gfp*, no social defeat stress: 19M, 14F), GFP-Post (AAV-*Gfp*, social defeat stress: 15M, 10F), SETD7-Pre (AAV-*Setd7* virus, no social defeat stress: 17M, 18F), SETD7-Post (AAV-*Setd7*, social defeat stress: 13M, 14F). Social interaction and open field exploration testing were performed as previously described (Balouek et al., 2023; Bennett et al., 2024; Peña et al., 2017). All behavior was recorded and initially analyzed with Ethovision software.

#### Social interaction

To assess social avoidance behavior, experimental mice were tested in a two-stage social interaction test. In the first 2.5-minute stage, experimental mice were placed into a novel empty arena (44 x 44 x 20 cm) with an empty enclosure along one wall. Mice were then removed, a novel aggressor mouse was placed into the enclosure in the “social interaction zone,” and experimental mice were reintroduced into the arena for an additional 2.5 minutes. Time spent in the interaction zones and in the corners was measured. Social interaction ratio (“SI ratio”) was calculated as time spent in the interaction zone with the aggressor present divided by time spent in the interaction zone without the aggressor present. “Susceptible” was defined as having a social interaction ratio <0.9, “resilient” as >1.1, and “indifferent” as in between.

#### Open field

Each mouse was allowed to explore an empty arena (44 x 44 x 20 cm) for 10 minutes and time spent in the center of the arena and overall distance traveled were measured.

#### Statistical analysis

Behavioral data were graphed and analyzed using Graphpad PRISM software (v. 10.0.3). Outliers were considered to be >2 standard-deviations away from group mean and were removed to avoid type-II error. Main effects and interactions were determined by mixed effects model to account for within-subject repeated testing, with stress, *Setd7*, sex, and interactions as fixed effects (type III). Šidák’s multiple comparisons testing was used for post hoc analyses including sex. Given a lack of significant main effect of sex, mixed-effects modeling was also run without sex as fixed effect, using Fisher’s LSD for post hoc analyses (reported in figures). Chi-squared contingency analysis was used to assess differences in proportion of resilient indifferent, and susceptible animals across groups. All significance thresholds were set at *p*<0.05; trends were reported for *p*<0.1. All statistical analyses are included in **Supplemental Table 3**.

### Nuclear isolation and ATAC-sequencing

To determine whether juvenile H3K4me1 enrichment by *Setd7* overexpression caused persistent changes in chromatin opening, juvenile mice (P9-12) received bilateral AAV-*Gfp* or AAV-*Setd7* as described above and were aged in control conditions until adulthood. Brains from control adult mice (P60-90) mice were harvested after cervical dislocation and rapid decapitation into ice-cold homogenization buffer (Corces et al., 2017). 1 mm slices of brain tissue were made using a slice matrix from which 14-gauge bilateral punches of the NAc were taken and flash frozen until use. Six GFP (3M, 3F) and five SETD7 (2M, 3F) samples from individual mice were used.

All nuclear isolation reactions were performed on ice. To isolate nuclei, bilateral NAc tissue punches from individual mice were dounce homogenized in 1.5uL of 1X Unstable homogenization buffer (Corces et al., 2017). Homogenate was transferred to 2 mL lo-bind tubes then spun down at 5000G for 5 min (at 4°C). The top 800uL of buffer was aspirated off and the sample was gently resuspended in remaining buffer. Nuclei were collected by iodixanol gradient and counts were collected using a hemocytometer. Yield of nuclei for all samples was between 3.5-5.8 million nuclei. DNA was then transposed using Illumina’s transposition kit (Illumina; cat#20034198) for 30 minutes at 37°C at 1,000RPM. Fragmented DNA was recovered with Qiagen MinElute kits (Qiagen; cat# 28604). Samples were amplified and purified according to (Corces et al., 2017) using Illumina UD indexes (Illumina; cat#20027214 A or B) and AMPure XP beads (ThermoFisher; cat#NC9959336). Resulting yield of DNA fragment sizes was determined by Bioanalyzer dsDNA High Sensitivity protocol and normalized by concentration for sequencing. Samples were pooled for sequencing on a single flowcell (NovaSeq S1 100nt Flowcell v1.5) at the Genomics Core Facility of the Lewis-Sigler Institute at Princeton University.

### ATAC-seq processing and analysis

ATAC-seq reads adapters were trimmed using fastp (v0.23.4) with base correction enabled and aligned to mm39 using bowtie2 (v2.4.5) (“--no-unal --no-mixed --no-discordant”). Using samtools (v1.13), reads with MAPQ>30 were retained, followed by samtools collate, fixmate, sort, and markdup to identify PCR duplicates. MACS2 (v2.2.9.1) was used to call narrowpeaks for each sample.

Quality control was assessed using Fastqc (v0.11.9), Picard CollectMultipleMetrics (v2.27.5), and multiqc (v1.17). We found read duplication rates between 33-54%, unique alignment rates between 78-86%, and final average deduplicated library size of 96M reads. We verified that mono- and di-nucleosome sized fragments were visible in all aligned insert size distributions. Using a custom R (v4.4.2) script and package rtracklayer (v1.66.0), we calculated a median of 41% for fraction of reads in MACS2 peaks.

A consensus peak set was generated using GenomicRanges (v1.58.0) “reduce” command in R. ATAC reads were imported using BRGenomics (v1.17.1) “import_bam” with “field=NULL” and unique reads in peaks were counted using “getCountsByRegions”. Differential accessibility was assessed using DESeq2 (v1.46.0) using a likelihood ratio test comparing a full model “~sex + condition” (with condition being SETD7 or GFP) to a reduced model of sex alone. Related plots were generated using ggplot2 (v3.5.1). We identified differentially accessible regions (DARs) as peaks with |log2FoldChange| >0.5 and p.adj<0.1.

ChIPseeker (v.1.36.0) was used to find genomic features overlapping differentially accessible regions (DARs), and to nominate putative gene contacts by taking the list of closest genes for the 43 most open regions (p.adj<0.1) and running gene ontology (GO) analysis. We used geneontology.org to perform GO term enrichment analysis on the putatively gene contacts of DARs.

To visualize tracks, RPM/CPM-normalized bigWig files were also generated in R using GenomicRanges “coverage” via BRGenomics “getStrandedCoverage” with strand information removed, and exported with rtracklayer.

### Patch-clamp electrophysiology

Male and female *Drd1a-tdTomato* mice were stereotactically injected in the NAc with AAV-*Gfp* or AAV-*Setd7* between P9-P12 as described above. Afterwards, animals were returned to the home cage, weaned at P21, and aged to adulthood in standard group-housing conditions. At >P60, female mice underwent 3 days of mild variable stress and were subsequently patched on the fourth day. The mild variable stress consisted of tail hang for one hour on the first day, tube restriction for one hour on the second day, and foot shock for one hour (100 random mild foot shocks at 0.45 mA) on the third day (Peña et al., 2019). At >P60, male mice were subjected to 10 days of subthreshold adult social defeat stress as described above and were patched on the eleventh day.

Mice were deeply anesthetized with isoflurane, rapidly decapitated, and brains removed into ice-cold choline cutting solution (composition in mM: 0.5 CaCl2, 110 C5H14CINO, 25 C6H12O6, 25 NaHCO3, 7 MgCl2, 11.6 C6H8O6, 3.1 C3H3NaO3, 2.5 KCl and 1.25 NaH2PO4). 210um coronal sections were made through the NAc on a vibratome (Leica VT 2100) and allowed to recover in carboxygenated artificial cerebrospinal fluid (aCSF; composition in mM: 119 NaCl, 2.5 KCl, 1.3 MgCl2, 2.5 CaCl2, 1.0 Na2HPO4, 26.2 NaHCO3, and 11 glucose) at 32°C for 15 minutes. Slices were then maintained at room temperature in aCSF until recording.

For whole cell patch clamp recordings, hemisected slices were superfused with aCSF containing 100 uM picrotoxin and 2 mM kynurenic acid at 30 ± 2°C in the recording chamber. D1-MSNs were identified by fluorescence, while non-fluorescent cells were recorded as putative D2-MSNs. For putative D2-MSNs, we avoided cells with large soma sizes and excluded any neurons that exhibited spontaneous firing or burst firing in response to current injection, consistent with striatal interneurons (Johansson and Silberberg, 2020; Kawaguchi, 1993).

Glass recording pipettes were pulled to a tip resistance of 3 - 5 MΩ and filled with a potassium gluconate-based internal solution (composition in mM: 130 potassium gluconate, 10 phosphocreatine disodium salt, 4 MgCl_2_, 3.4 Na_2_ATP, 0.1 Na_3_GTP, 1.1 EGTA, 5 HEPES). All recordings were made in whole cell configuration. To generate an input-output curve of neuronal excitability, 600 ms current steps were injected in increments of 25 pA from −150 to 500 pA and the numbers of spikes were quantified for each step. The current step at which the cell fired the first action potential was determined as the rheobase. Voltage sag was also measured from the same recordings using the negative current steps. Input resistance was calculated based on the average voltage change in response to 20 successive −10 pA current injections. Recordings were excluded if the series resistance varied by more than 20% over the course of recordings. Patch-clamp electrophysiology data acquisition was completed using Clampex 11.4 software (Molecular Devices).

### Histone modification analysis

Analysis of post-translational histone modifications in NAc across three developmental time points (P21, P35, P70-90, n=3 males/group/age) was done by re-analyzing previously published histone post-translational modification mass spectrometry (data from (Kronman et al., 2021). This data represents percentage of each measured histone modification per fragmented histone peptide (for example, percentage of unmodified histone-3 lysine-4, H3K4me1, H3K4me2, and H3K4me3). Data for H3K27me2 and H3K27me3 are summations of the modification of interest alone or in combination with any H3K36 modification measured on the same peptide fragment. For each histone modification of interest, two-way ANOVA was done to assess main effects and interactions of ELS with age.

### Immunohistochemistry and imaging

To confirm viral targeting of the NAc following behavioral experiments, a subset of mice was anesthetized by i.p. injection of 100 mg/kg ketamine and 10 mg/kg xylazine, then transcardially perfused with ice-cold saline followed by 4% paraformaldehyde (PFA). Brains were dissected and submerged in fresh 4% PFA at 4°C overnight, then stored in 30% sucrose solution for an additional 2 days (until the tissue became isotonic). Brains were then kept at −80°C until slicing. For slicing, brains were embedded in Tissue-Plus™ O.C.T. Compound (Fisher Scientific) and sliced in 20µm sections on a cryostat at −20°C. Slices were mounted directly onto Superfrost™ Plus microscope slides (Fisher Scientific) and stored at −20°C until staining.

Immunohistochemistry entailed the following steps (at room temperatures unless noted): (1) Rinsing the slices in 1 X phosphate-buffered saline (PBS) 3 times for 5 min each. (2) Incubating the slices in 1.5% normal donkey serum (NSD) in 1 X PBS-Triton X-100 solution for 1 h. (3) Incubating the slices in primary antibody [1:500 Anti-Green Fluorescent Protein (Chicken) Antibody (Aves Labs); 1:150 SETD7 Mouse Monoclonal Antibody (OriGene); and 1.5% NSD in 1 X PBS-Triton X-100 solution] overnight at 4°C. (4) Rinsing the slices in 1 X PBS 3 times for 5 min each. (5) Incubating the slices in a secondary antibody solution [1:500 Alexa Fluor® 488 AffiniPure Donkey Anti-Chicken IgY (IgG) (H+L) (Jackson ImmunoReseach Labs); 1:500 Cy™3 AffiniPure Donkey Anti-Mouse IgG (H+L) (Jackson ImmunoReseach Labs); and 1.5% NSD in 1 X PBS-Triton X-100] for 1 h. (6) Rinsing the slices in 1 X PBS 3 times for 5 min each. (7) Incubating the slices in 0.0025% DAPI solution for 5 min. (8) Rinsing the slices in 1 X PBS 3 times for 5 min each. Following the last rinse, Fluoromount-G® mounting media (SouthernBiotech) was used to coverslip the immune-stained slices. Samples were kept at 4°C until imaging. Fluorescent imaging was performed with a Nanozoomer S60 slide scanner (Hamamatsu). All images were processed using the ImageJ (version 1.2.0) software containing the FIJI plugin (NIH, Bethesda, MD, USA).

## Data Availability

All sequencing data is deposited in NCBI’s Gene Expression Omnibus (GEO), accession number GSE268468. Sequencing data was analyzed via standardized pipelines as described.

## SUPPLEMENTAL MATERIAL

***Supplemental Figure S1:*** Additional post-translational histone modifications across development in male NAc

***Supplemental Figure S2:*** Juvenile *Setd7* overexpression in NAc does not significantly alter D1-MSN currents

***Supplemental Figure S3:*** Adult *Setd7* overexpression in NAc minimally influences stress response

***Supplemental Table 1:*** ATAC-seq differentially accessible region analysis

***Supplemental Table 2:*** DAR-primed genes

***Supplemental Table 3:*** Statistical analysis of behavior after juvenile or adult *Setd7* overexpression

## Notes

### Competing Interest Statement

The authors have declared no competing interest.

### Summary of Updates

This revised manuscript now focuses on the finding of increased H3K4me1 in NAc and subsequent Setd7 manipulations, including addition of new electrophysiological recordings after Setd7 manipulations and additional behavior. Previously included experience-tagged cell ATAC-seq was removed after difficulty replicating using snRNA-seq; instead we now present ATAC-seq after juvenile Setd7 overexpression. Supplemental files are updated. The authors have been updated to reflect these changes and new data. While these edits and additions are comprehensive, they focus the manuscript and extend mechanistic investigation.

## REFERENCES

Balouek J-A, Mclain CA, Minerva AR, Rashford RL, Bennett SN, Rogers FD, Peña CJ (2023) Reactivation of Early-Life Stress-Sensitive Neuronal Ensembles Contributes to Lifelong Stress Hypersensitivity. J Neurosci 43:5996–6009.

Bariselli S, Fobbs WC, Creed MC, Kravitz AV (2019) A competitive model for striatal action selection. Brain Research 1713:70–79.

Bastle RM, Maze I (2019) Chromatin Regulation in Complex Brain Disorders. Current opinion in behavioral sciences 25:57–65.

Bath KG, Manzano-Nieves G, Goodwill H (2016) Early life stress accelerates behavioral and neural maturation of the hippocampus in male mice. Hormones and Behavior 82:64– 71.

Bennett SN, Chang AB, Rogers FD, Jones P, Peña CJ (2024) Thyroid hormones mediate the impact of early-life stress on ventral tegmental area gene expression and behavior. Hormones and Behavior 159:105472.

Bonifer C, Cockerill PN (2017) Chromatin priming of genes in development: Concepts, mechanisms and consequences. Experimental Hematology 49:1–8.

Burns SB, Szyszkowicz JK, Luheshi GN, Lutz P-E, Turecki G (2018) Plasticity of the epigenome during early-life stress. Seminars in Cell & Developmental Biology 77:115–132.

Calo E, Wysocka J (2013) Modification of Enhancer Chromatin: What, How, and Why? Molecular Cell 49:825–837.

Corces MR et al. (2017) An improved ATAC-seq protocol reduces background and enables interrogation of frozen tissues. Nature Methods 14:959–962.

Covington HE, Maze I, Sun H, Bomze HM, DeMaio KD, Wu EY, Dietz DM, Lobo MK, Ghose S, Mouzon E, Neve RL, Tamminga CA, Nestler EJ (2011) A role for repressive histone methylation in cocaine-induced vulnerability to stress. Neuron 71:656–670.

Creed M (2018) Current and emerging neuromodulation therapies for addiction: insight from pre-clinical studies. Current Opinion in Neurobiology 49:168–174.

Creed M, Ntamati NR, Chandra R, Lobo MK, Lüscher C (2016) Convergence of Reinforcing and Anhedonic Cocaine Effects in the Ventral Pallidum. Neuron 92:214–226.

Creed M, Pascoli VJ, Lüscher C (2015) Refining deep brain stimulation to emulate optogenetic treatment of synaptic pathology. Science 347:659–664.

Dias C et al. (2014) β-catenin mediates stress resilience through Dicer1/microRNA regulation. Nature 516:51–55.

Dobbs LK, Kaplan AR, Lemos JC, Matsui A, Rubinstein M, Alvarez VA (2016) Dopamine Regulation of Lateral Inhibition between Striatal Neurons Gates the Stimulant Actions of Cocaine. Neuron 90:1100–1113.

Duque-Wilckens N, Torres LY, Yokoyama S, Minie VA, Tran AM, Petkova SP, Hao R, Ramos-Maciel S, Rios RA, Jackson K, Flores-Ramirez FJ, Garcia-Carachure I, Pesavento PA, Iñiguez SD, Grinevich V, Trainor BC (2020) Extrahypothalamic oxytocin neurons drive stress-induced social vigilance and avoidance. Proceedings of the National Academy of Sciences USA:202011890.

Fang LZ, Creed MC (2024) Updating the striatal–pallidal wiring diagram. Nat Neurosci 27:15–27.

Farrelly LA et al. (2019) Histone serotonylation is a permissive modification that enhances TFIID binding to H3K4me3. Nature 128:1.

Ferrari KJ, Scelfo A, Jammula S, Cuomo A, Barozzi I, Stützer A, Fischle W, Bonaldi T, Pasini D (2014) Polycomb-Dependent H3K27me1 and H3K27me2 Regulate Active Transcription and Enhancer Fidelity. Molecular Cell 53:49–62.

Francis TC, Chandra R, Friend DM, Finkel E, Dayrit G, Miranda J, Brooks JM, Iñiguez SD, O’Donnell P, Kravitz A, Lobo MK (2014) Nucleus Accumbens Medium Spiny Neuron Subtypes Mediate Depression-Related Outcomes to Social Defeat Stress. Biological Psychiatry.

Francis TC, Lobo MK (2016) Emerging Role for Nucleus Accumbens Medium Spiny Neuron Subtypes in Depression. Biological Psychiatry Available at: http://linkinghub.elsevier.com/retrieve/pii/S0006322316328098.

Franklin A, Davies JP, Clifton NE, Blake GET, Bamford R, Walker EM, Chioza B, Frith M, Burrage J, Owens N, Prabhakar S, Dempster E, Hannon E, Mill J (2025) Cell-type-specific DNA methylation dynamics in the prenatal and postnatal human cortex. bioRxiv Available at: https://www.biorxiv.org/content/10.1101/2025.02.21.639467v1 [Accessed April 7, 2025].

Gee D, Gabard-Durnam LJ, Flannery J, Goff B, Humphreys KL, Telzer EH, Hare TA, Bookheimer SY, Tottenham N (2013) Early developmental emergence of human amygdala– prefrontal connectivity after maternal deprivation. Proceedings of the National Academy of Sciences 110:15638– 15643.

Geiger LT, Balouek J-A, Barrett MR, Thompson JM, Fang LZ, Farrelly LA, Chen AS, Tang M, Bennett SN, Garcia BA, Maze I, Creed MC, Peña CJ (2024) Early-life stress alters chromatin modifications in VTA to prime stress sensitivity. bioRxiv Available at: http://biorxiv.org/lookup/doi/10.1101/2024.03.14.584631 [Accessed May 7, 2025].

Golden SA, Covington HE, Berton O, Russo SJ (2011) A standardized protocol for repeated social defeat stress in mice. Nature Protocols 6:1183–1191.

Grant CE, Bailey TL (2021) XSTREME: Comprehensive motif analysis of biological sequence datasets. bioRxiv Available at: https://www.biorxiv.org/content/10.1101/2021.09.02.458722v1 [Accessed April 9, 2024].

Griffith EC, West AE, Greenberg ME (2024) Neuronal enhancers fine-tune adaptive circuit plasticity. Neuron 112:3043–3057.

Hanson JL, Williams AV, Bangasser DA, Peña CJ (2021) Impact of Early Life Stress on Reward Circuit Function and Regulation. Frontiers in Psychiatry 12:1799.

Heffel MG et al. (2024) Temporally distinct 3D multi-omic dynamics in the developing human brain. Nature 635:481– 489.

Heintzman ND, Stuart RK, Hon G, Fu Y, Ching CW, Hawkins RD, Barrera LO, Van Calcar S, Qu C, Ching KA, Wang W, Weng Z, Green RD, Crawford GE, Ren B (2007) Distinct and predictive chromatin signatures of transcriptional promoters and enhancers in the human genome. Nat Genet 39:311–318.

Heller EA et al. (2014) Locus-specific epigenetic remodeling controls addiction- and depression-related behaviors. Nature Neuroscience 17:1720–1727.

Herring CA et al. (2022) Human prefrontal cortex gene regulatory dynamics from gestation to adulthood at single-cell resolution. Cell 185:4428-4447.e28.

Hodes GE et al. (2015) Sex Differences in Nucleus Accumbens Transcriptome Profiles Associated with Susceptibility versus Resilience to Subchronic Variable Stress. Journal of Neuroscience 35:16362–16376.

Howerton CL, Morgan CP, Fischer DB, Bale TL (2013) O-GlcNAc transferase (OGT) as a placental biomarker of maternal stress and reprogramming of CNS gene transcription in development. Proceedings of the National Academy of Sciences 110:5169–5174.

Ianov L, Day JJ (2024) Ratlas. Available at: https://zenodo.org/records/10967245 [Accessed June 23, 2025].

Johansson Y, Silberberg G (2020) The Functional Organization of Cortical and Thalamic Inputs onto Five Types of Striatal Neurons Is Determined by Source and Target Cell Identities. Cell Reports 30:1178-1194.e3.

Kawaguchi Y (1993) Physiological, morphological, and histochemical characterization of three classes of interneurons in rat neostriatum. J Neurosci 13:4908–4923.

Kofent J, Zhang J, Spagnoli FM (2016) The histone methyltransferase Setd7 promotes pancreatic progenitor identity. Development 143:3573–3581.

Kos A, Lopez JP, Bordes J, de Donno C, Dine J, Brivio E, Karamihalev S, Luecken MD, Almeida-Correa S, Gasperoni S, Dick A, Miranda L, Büttner M, Stoffel R, Flachskamm C, Theis FJ, Schmidt MV, Chen A (2023) Early life adversity shapes social subordination and cell type–specific transcriptomic patterning in the ventral hippocampus. Science Advances 9:eadj3793.

Kravitz AV, Tye LD, Kreitzer AC (2012) Distinct roles for direct and indirect pathway striatal neurons in reinforcement. Nat Neurosci 15:816–818.

Kronman H, Torres-Berrío A, Sidoli S, Issler O, Godino A, Ramakrishnan A, Mews P, Lardner CK, Parise EM, Walker DM, van der Zee YY, Browne CJ, Boyce BF, Neve R, Garcia BA, Shen L, Peña CJ, Nestler EJ (2021) Long-term behavioral and cell-type-specific molecular effects of early life stress are mediated by H3K79me2 dynamics in medium spiny neurons. Nature Neuroscience:1–10.

Labonté B, Suderman M, Maussion G, Navaro L, Yerko V, Mahar I, Bureau A, Mechawar N, Szyf M, Meaney MJ, Turecki G (2012) Genome-wide epigenetic regulation by early-life trauma. Archives of general psychiatry 69:722–731.

Lemos JC, Friend DM, Kaplan AR, Shin JH, Rubinstein M, Kravitz AV, Alvarez VA (2016) Enhanced GABA Transmission Drives Bradykinesia Following Loss of Dopamine D2 Receptor Signaling. Neuron 90:824–838.

Lepack AE et al. (2020) Dopaminylation of histone H3 in ventral tegmental area regulates cocaine seeking. Science 368:197– 201.

Lepack AE, Bagot RC, Peña CJ, Loh Y-HE, Farrelly LA, Lu Y, Powell SK, Lorsch ZS, Issler O, Cates HM, Tamminga CA, Molina H, Shen L, Nestler EJ, Allis CD, Maze I (2016) Aberrant H3.3 dynamics in NAc promote vulnerability to depressive-like behavior. Proceedings of the National Academy of Sciences USA 113:12562–12567.

Levine A, Worrell TR, Zimnisky R, Schmauss C (2012) Early life stress triggers sustained changes in histone deacetylase expression and histone H4 modifications that alter responsiveness to adolescent antidepressant treatment. Neurobiology of disease 45:488–498.

Lister R et al. (2013) Global Epigenomic Reconfiguration During Mammalian Brain Development. Science 341:1237905– 1237905.

Lobo MK, Covington HE, Chaudhury D, Friedman AK, Sun H, Damez-Werno D, Dietz DM, Zaman S, Koo JW, Kennedy PJ, Mouzon E, Mogri M, Neve RL, Deisseroth K, Han M-H, Nestler EJ (2010) Cell type-specific loss of BDNF signaling mimics optogenetic control of cocaine reward. Science 330:385–390.

Long HK, Prescott SL, Wysocka J (2016) Ever-Changing Landscapes: Transcriptional Enhancers in Development and Evolution. Cell 167:1170–1187.

Lutz P-E et al. (2017) Association of a History of Child Abuse With Impaired Myelination in the Anterior Cingulate Cortex: Convergent Epigenetic, Transcriptional, and Morphological Evidence. AJP 174:1185–1194.

Lutz P-E, Chay M-A, Pacis A, Chen GG, Aouabed Z, Maffioletti E, Théroux J-F, Grenier J-C, Yang J, Aguirre M, Ernst C, Redensek A, van Kempen LC, Yalcin I, Kwan T, Mechawar N, Pastinen T, Turecki G (2021) Non-CG methylation and multiple histone profiles associate child abuse with immune and small GTPase dysregulation. Nat Commun 12:1132.

Matikainen-Ankney BA, Legaria AA, Pan Y, Vachez YM, Murphy CA, Schaefer RF, McGrath QJ, Wang JG, Bluitt MN, Ankney KC, Norris AJ, Creed MC, Kravitz AV (2023) Nucleus Accumbens D1 Receptor–Expressing Spiny Projection Neurons Control Food Motivation and Obesity. Biological Psychiatry 93:512–523.

Mercer EM, Lin YC, Benner C, Jhunjhunwala S, Dutkowski J, Flores M, Sigvardsson M, Ideker T, Glass CK, Murre C (2011) Multilineage priming of enhancer repertoires precedes commitment to the B and myeloid cell lineages in hematopoietic progenitors. Immunity 35:413–425.

Morrison KE, Cole AB, Kane PJ, Meadows VE, Thompson SM, Bale TL (2020) Pubertal adversity alters chromatin dynamics and stress circuitry in the pregnant brain. Neuropsychopharmacology 69:1–9.

Muir J, Tse YC, Iyer ES, Biris J, Cvetkovska V, Lopez J, Bagot RC (2020) Ventral Hippocampal Afferents to Nucleus Accumbens Encode Both Latent Vulnerability and Stress-Induced Susceptibility. Biological Psychiatry 3223(20)31627–9.

Parade SH, Huffhines L, Daniels TE, Stroud LR, Nugent NR, Tyrka AR (2021) A systematic review of childhood maltreatment and DNA methylation: candidate gene and epigenome-wide approaches. Translational Psychiatry 11:1–33.

Parel ST, Bennett SN, Cheng CJ, Timmermans OC, Fiori LM, Turecki G, Peña CJ (2023) Transcriptional signatures of early-life stress and antidepressant treatment efficacy. Proceedings of the National Academy of Sciences 120:e2305776120.

Parel ST, Peña CJ (2020) Genome-wide signatures of early life stress: influence of sex. Biological Psychiatry 3223(20)32119–3.

Pascoli V, Terrier J, Espallergues J, Valjent E, O’Connor EC, Lüscher C (2014) Contrasting forms of cocaine-evoked plasticity control components of relapse. Nature 509:459– 464.

Peña CJ (2017) D1 and D2 Type Medium Spiny Neuron Contributions to Depression. Biological Psychiatry 81:636– 638.

Peña CJ (2025) Epigenetic regulation of brain development, plasticity, and response to early-life stress. Neuropsychopharmacol:1–11.

Peña CJ, Kronman HG, Walker DM, Cates HM, Bagot RC, Purushothaman I, Issler O, Loh Y-HE, Leong T, Kiraly DD, Goodman E, Neve RL, Shen L, Nestler EJ (2017) Early life stress confers lifelong stress susceptibility in mice via ventral tegmental area OTX2. Science 356:1185–1188.

Peña CJ, Neugut YD, Champagne FA (2013) Developmental Timing of the Effects of Maternal Care on Gene Expression and Epigenetic Regulation of Hormone Receptor Levels in Female Rats. Endocrinology 154:4340–4351.

Peña CJ, Smith M, Ramakrishnan A, Cates HM, Bagot RC, Kronman HG, Patel B, Chang AB, Purushothaman I, Dudley J, Morishita H, Shen L, Nestler EJ (2019) Early life stress alters transcriptomic patterning across reward circuitry in male and female mice. Nature Communications 10:5098.

Phillips RA, Tuscher JJ, Fitzgerald ND, Wan E, Zipperly ME, Duke CG, Ianov L, Day JJ (2023) Distinct subpopulations of D1 medium spiny neurons exhibit unique transcriptional responsiveness to cocaine. Molecular and Cellular Neuroscience 125:103849.

Pignatelli M, Tejeda HA, Barker DJ, Bontempi L, Wu J, Lopez A, Palma Ribeiro S, Lucantonio F, Parise EM, Torres-Berrio A, Alvarez-Bagnarol Y, Marino RAM, Cai Z-L, Xue M, Morales M, Tamminga CA, Nestler EJ, Bonci A (2021) Cooperative synaptic and intrinsic plasticity in a disynaptic limbic circuit drive stress-induced anhedonia and passive coping in mice. Mol Psychiatry 26:1860–1879.

Pusalkar M, Suri D, Kelkar A, Bhattacharya A, Galande S, Vaidya VA (2015) Early stress evokes dysregulation of histone modifiers in the medial prefrontal cortex across the life span. Developmental Psychobiology:n/a-n/a.

Rahman MF, McGowan PO (2022) Cell-type-specific epigenetic effects of early life stress on the brain. Transl Psychiatry 12:1–10.

Rakesh D, Whittle S, Sheridan MA, McLaughlin KA (2023) Childhood socioeconomic status and the pace of structural neurodevelopment: accelerated, delayed, or simply different? Trends in Cognitive Sciences https://www.sciencedirect.com/science/article/pii/S1364661323000736.

Rashford RL, DeBerardine M, Dan S, Bennett SN, Peña CJ (2025) Persistent open chromatin state in early-life stress-activated cells of the VTA. Scientific Reports 15:36118.

Reh RK, Dias BG, Nelson CA, Kaufer D, Werker JF, Kolb B, Levine JD, Hensch TK (2020) Critical period regulation across multiple timescales. Proceedings of the National Academy of Sciences USA:201820836.

Savolainen MH, Richie CT, Harvey BK, Männistö PT, Maguire-Zeiss KA, Myöhänen TT (2014) The beneficial effect of a prolyl oligopeptidase inhibitor, KYP-2047, on alpha-synuclein clearance and autophagy in A30P transgenic mouse. Neurobiology of Disease 68:1–15.

Schoenfelder S, Fraser P (2019) Long-range enhancer–promoter contacts in gene expression control. Nat Rev Genet 20:437– 455.

Short AK, Thai CW, Chen Y, Kamei N, Pham AL, Birnie MT, Bolton JL, Mortazavi A, Baram TZ (2023) Single-Cell Transcriptional Changes in Hypothalamic Corticotropin-Releasing Factor–Expressing Neurons After Early-Life Adversity Inform Enduring Alterations in Vulnerabilities to Stress. Biological Psychiatry Global Open Science 3:99–109.

Simmons RK, Stringfellow SA, Glover ME, Wagle AA, Clinton SM (2013) DNA methylation markers in the postnatal developing rat brain. Brain Research 1533:26–36.

Stroud H, Su SC, Hrvatin S, Greben AW, Renthal W, Boxer LD, Nagy MA, Hochbaum DR, Kinde B, Gabel HW, Greenberg ME (2017) Early-Life Gene Expression in Neurons Modulates Lasting Epigenetic States. Cell 171:1151-1164.e16.

Stroud H, Yang MG, Tsitohay YN, Davis CP, Sherman MA, Hrvatin S, Ling E, Greenberg ME (2020) An Activity-Mediated Transition in Transcription in Early Postnatal Neurons. Neuron 0896–6273.

Sun H et al. (2015) ACF chromatin-remodeling complex mediates stress-induced depressive-like behavior. Nature medicine 21:1146–1153.

Torres-Berrío A, Estill M, Patel V, Ramakrishnan A, Kronman H, Minier-Toribio A, Issler O, Browne CJ, Parise EM, van der Zee YY, Walker DM, Martínez-Rivera FJ, Lardner CK, Durand-de Cuttoli R, Russo SJ, Shen L, Sidoli S, Nestler EJ (2024) Mono-methylation of lysine 27 at histone 3 confers lifelong susceptibility to stress. Neuron 112:2973-2989.e10.

Weaver ICG, Cervoni N, Champagne FA, D’Alessio AC, Sharma S, Seckl JR, Dymov S, Szyf M, Meaney MJ (2004) Epigenetic programming by maternal behavior. Nature Neuroscience 7:847–854.

Weder N, Zhang H, Jensen K, Yang B-Z, Simen A, Jackowski A, Lipschitz D, Douglas-Palumberi H, Ge M, Perepletchikova F, O’Loughlin K, Hudziak JJ, Gelernter J, Kaufman J (2014) Child Abuse, Depression, and Methylation in Genes Involved With Stress, Neural Plasticity, and Brain Circuitry. Journal of the American Academy of Child and Adolescent Psychiatry 53:417-424.e5.

Yohn C, Dieterich A, Bazer A, Maita I, Giedraitis M, Samuels B (2019) Chronic Non-Discriminatory Social Defeat Is an Effective Chronic Stress Paradigm for Both Male and Female Mice. Neuropsychopharmacology 44:2220–2229.

