## Supplemental Figures 1-3 for "Altered postnatal chromatin development in the nucleus accumbens primes lifelong stress sensitivity"

*for*

***Supplemental Figure S1:*** Additional post-translational histone modifications across development in male NAc

***Supplemental Figure S2:*** Juvenile *Setd7* overexpression in NAc does not significantly alter D1-MSN currents

***Supplemental Figure S3:*** Adult *Setd7* overexpression in NAc minimally influences stress response

***Supplemental Table 1:*** ATAC-seq differentially accessible region analysis

***Supplemental Table 2:*** DAR-primed genes

***Supplemental Table 3:*** Statistical analysis of behavior after juvenile or adult *Setd7* overexpression

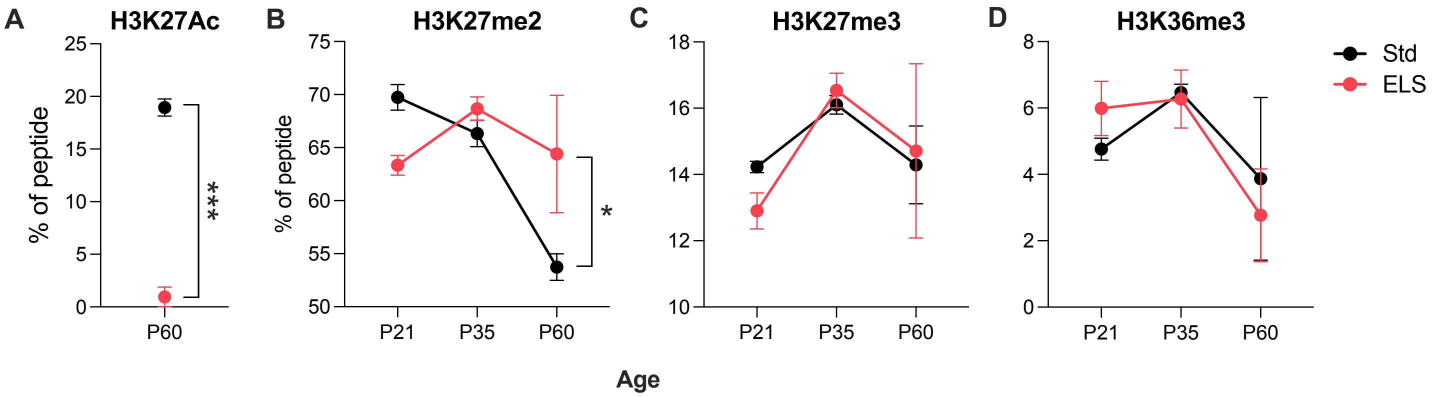


**Supplemental Figure S1: Additional post-translational histone modifications across development in male NAc.** Re-analysis and visualization of mass-spectrometry data published in Kronman et al, *Nature Neuroscience*, 2021. (A) H3K27Ac was undetectable at P21 and P35. P60 Welch's two-tailed t-test: $t(1,5)=14.50$, $p=0.0002$. (B) H3K27me2: interaction between ELS and age: $F(1, 12)=5.874$, $p=0.017$. (C) H3K27me3, not significant. (D) H3K36me3, not significant. All data depicted as mean±SEM.

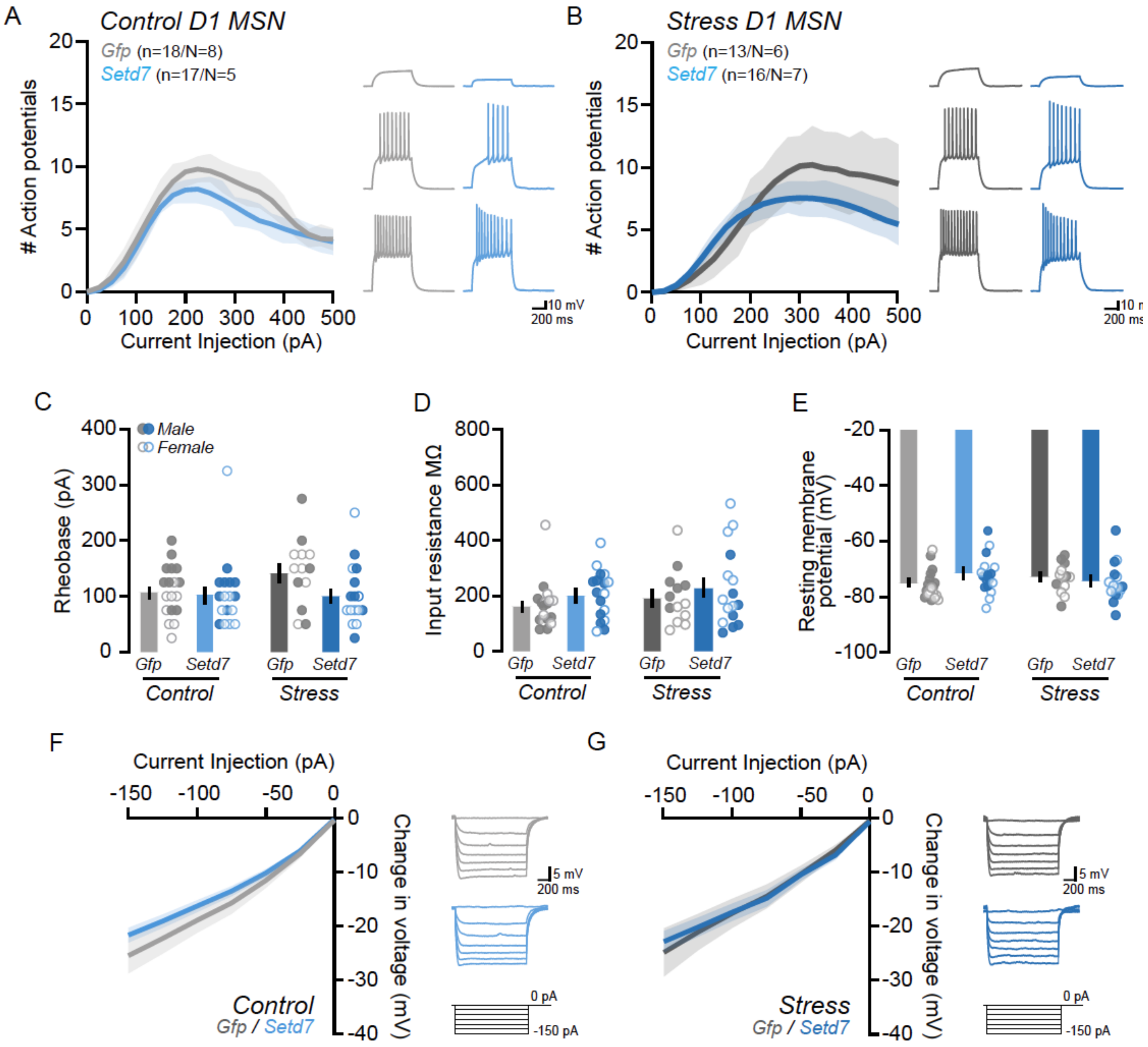


**Supplemental Figure S2**: **Juvenile *Setd7* overexpression in NAc does not significantly alter D1-MSN currents.**

**A-B.** (Left) Input-output plot of action potentials elicited in response to successive 600ms driving currents from control (**A**) and stressed (**B**) mice. (Right) Representative traces depict action potential firing in response to a 50 pA (top), 150 pA (middle), and 250 pA (bottom) driving current. *Setd7* overexpression did not affect action potential firing in D1 MSNs under control (F(1,33)=1.473, *p*=0.2335, two-way RM ANOVA) or stress (F(1,27)=0.3265, *p*=0.5725, two-way RM ANOVA) conditions.

**C-E**. The effect of *Setd7* overexpression on intrinsic membrane properties of D1 MSNs. *Setd7* overexpression does not alter the rheobase (**C**; Control: *p*=0.4114, *U*=128.0; Stress: *p*=0.0634, *U*=62.00, Mann-Whitney test), input resistance (**D**; Control: *p*=0.1028, *U*=103.0; Stress: *p*=0.8123, *U*=98.00, Mann-Whitney test), nor the resting membrane potential (**E**; Control: *p*=0.0537, *U*=94.50; Stress: *p*=0.2824, *U*=79.00, Mann-Whitney test) of D1 MSNs in control or stressed mice.

**F-G**. The peak voltage change in response to successive 600ms hyperpolarizing currents from control (**F**) and stressed (**G**) mice. *Setd7* overexpression does not affect the peak voltage sag under control (F(1,33)=1.450, *p*=0.2370, two-way RM ANOVA) or stress (F(1,27)=0.1954, *p*=0.6619, two-way RM ANOVA) conditions. All data are depicted as mean ±SEM.

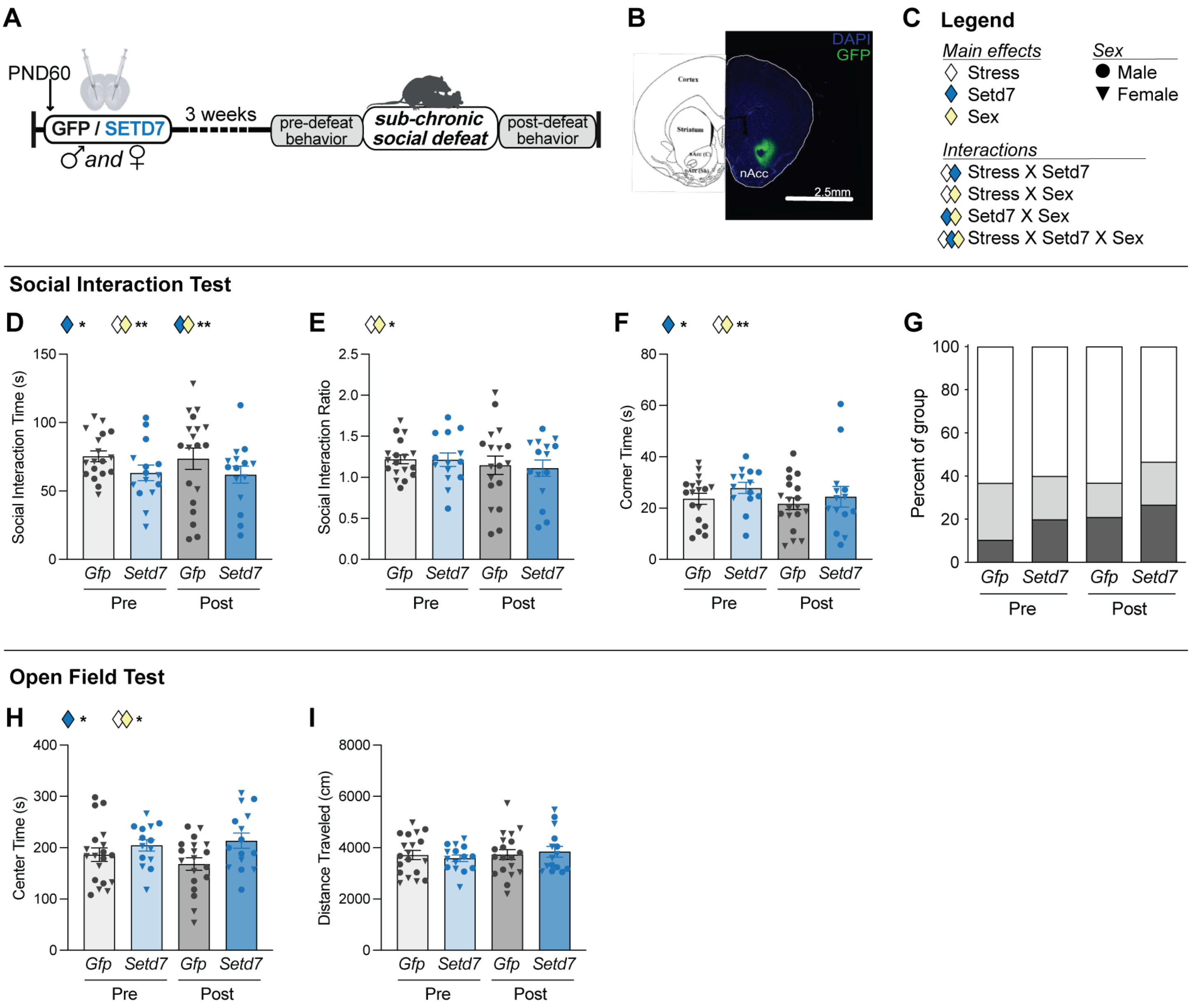


**Supplemental Figure S3: *Setd7* overexpression in adult NAc minimally influences stress response**.
(A) Experimental Timeline. (B) Representative image and brain atlas showing adult viral targeting of NAc. Scale bar, 2.5mm. (C) Key for statistical main effects and interactions (not all effects/interactions found). Within the social interaction test, (D) social interaction time, (E) social interaction ratio, (F)time spent in the corners, and (G) percent of each group considered resilient (open), indifferent (medium gray), or susceptible (dark gray).
Within the open field test, (H) time spent in the center of the arena and (I) total distance traveled before (pre) and after (post) sub-chronic social defeat stress.
Main effects and interactions from mixed-effect modeling are indicated above each plot. ***p*<0.01; **p*<0.05. Data are represented as mean ± SEM.
